# A tRNA-specific function for tRNA methyltransferase Trm10 is associated with a new tRNA quality control mechanism in *Saccharomyces cerevisiae*

**DOI:** 10.1101/2023.10.06.561306

**Authors:** Isobel E. Bowles, Jane E. Jackman

## Abstract

In *Saccharomyces cerevisiae* a single homolog of the tRNA methyltransferase Trm10 performs m^1^G9 modification on 13 different tRNAs. Here we provide evidence that the m^1^G9 modification catalyzed by *S. cerevisiae* Trm10 plays a biologically important role for one of these tRNA substrates, tRNA^Trp^. Overexpression of tRNA^Trp^ (and not any of 38 other elongator tRNAs) rescues growth hypersensitivity of the *trm10Δ* strain in the presence of the antitumor drug 5-fluorouracil (5FU). Mature tRNA^Trp^ is depleted in *trm10Δ* cells, and its levels are further decreased upon growth in 5FU, while another Trm10 substrate (tRNA^Gly^) is not affected under these conditions. Thus, m^1^G9 in *S. cerevisiae* is another example of a tRNA modification that is present on multiple tRNAs but is only essential for the biological function of one of those species. In addition to the effects of m^1^G9 on mature tRNA^Trp^, precursor tRNA^Trp^ species accumulate in the same strains, an effect that is due to at least two distinct mechanisms. The levels of mature tRNA^Trp^ are rescued in the *trm10Δmet22Δ* strain, consistent with the known role of Met22 in tRNA quality control, where deletion of *met22* causes inhibition of 5’-3’ exonucleases that catalyze tRNA decay. However, none of the known Met22-associated exonucleases appear to be responsible for decay of hypomodified tRNA^Trp^, based on inability of mutants of each enzyme to rescue growth of the *trm10Δ* strain in the presence of 5FU. Thus, the surveillance of tRNA^Trp^ appears to constitute a distinct tRNA quality control pathway in *S. cerevisiae*.

## INTRODUCTION

Transfer ribonucleic acid (tRNA) molecules are the most highly modified RNA molecules in the cell. Cytoplasmic tRNAs in *Saccharomyces cerevisiae* have known modifications at 36 of the ∼75 nucleotide positions and each tRNA has an average of 12.6 modifications (Juhling et al. 2009; Machnicka et al. 2013; Phizicky and Hopper 2023). tRNA modifications to the anticodon stem-loop (ACL) are often performed by genes that, when deleted, exhibit translation-related phenotypic defects due to their important roles in the efficiency or fidelity of translation (Phizicky and Hopper 2023). tRNA modifications found in the body of the tRNA, however, are performed by enzymes that often display mild to no obvious phenotypes when their genes are deleted in model systems such as *S. cerevisiae*. Recently, many of these genes have been implicated in more subtle roles that are often specific to unique tRNA species (Phizicky and Alfonzo 2010; Jackman and Alfonzo 2013; Howell and Jackman 2019; Phizicky and Hopper 2023). In cases studied so far, tRNA body modifications are generally thought to improve overall tRNA folding and stability and thus aid the tRNA in evading degradation by quality control pathways that remove low-quality tRNA from the pool of molecules available for translation (Whipple et al. 2011; Hopper and Huang 2015; Phizicky and Hopper 2023).

Multiple tRNA decay pathways in *S. cerevisiae* act to degrade specific tRNA species upon the loss of certain tRNA modifications or in the case of aberrant processing (Kadaba et al. 2004; Alexandrov et al. 2006; Hopper and Huang 2015; Payea et al. 2020; Tasak and Phizicky 2022). If the tRNA does not maintain its correct structure, including due to improper modification, or is lingering in the nucleus due to improper processing or trafficking, the tRNA is probably not sufficiently fit for its role in translation, and its removal helps to maintain the overall efficiency of protein synthesis. The tRNA quality control pathways so far identified in *S. cerevisiae* include the rapid tRNA decay (RTD) pathway that acts on mature hypomodified tRNA in both the nucleus and cytoplasm (Alexandrov et al. 2006; Chernyakov et al. 2008), the Met22-dependent decay pathway (MPD) which targets pre-tRNA with an aberrant intron-exon junction (Payea et al. 2020), and the nuclear TRAMP complex which acts on aberrant initiator or pre-tRNA, although RTD was also recently shown to contribute to surveillance of this tRNA (Kadaba et al. 2004; Tasak and Phizicky 2022). Interestingly, in all cases where RTD has been identified in connection with the loss of a tRNA modifying enzyme, only a subset of the tRNAs normally modified by the enzyme are substantially degraded upon the loss of the modification (Alexandrov et al. 2006; Kotelawala et al. 2008; Han et al. 2015). The stability of tRNA species varies and each tRNA requires different modifications to differing degrees, which is sensed distinctly by the exonucleases associated with each degradation pathway.

The tRNA methyltransferase 10 (Trm10) was first identified in *S. cerevisiae* as an m^1^G9 methyltransferase, but is conserved throughout Archaea and Eukarya (Jackman et al. 2003). Trm10 modifies the core of tRNA and homologs from some species have been identified that act as m^1^G9 and/or m^1^A9 methyltransferases, despite all of these enzymes sharing a similar protein fold as members of the SPOUT superfamily (Kempenaers et al. 2010; Vilardo et al. 2012; Howell et al. 2019; Vilardo et al. 2020; Strassler et al. 2022). Trm10 has three paralogs in humans, with TRMT10A acting as a m^1^G9 methyltransferase on multiple tRNAs and TRMT10B performing m^1^A9 on only tRNA^Asp^, while TrmT10C is a bifunctional m^1^A9/m^1^G9 methyltransferase that functions in human mitochondria as part of an unusual protein-only RNaseP complex (Holzmann et al. 2008; Vilardo et al. 2012; Howell et al. 2019; Vilardo et al. 2020). tRNA modification has proved increasingly important to human health (Torres et al. 2014; Suzuki 2021). Nine different homozygous loss of function mutations in *TRMT10A* have been identified in patients, and these are associated with a syndrome that primarily manifests in metabolic and neurological disorders in the affected patients (Igoillo-Esteve et al. 2013; Gillis et al. 2014; Narayanan et al. 2015; Zung et al. 2015; Yew et al. 2016; Lin et al. 2020; Siklar et al. 2021; Stern et al. 2022). Interestingly, for one of these patients, having a premature stop codon mutation of *TRMT10A* leads human TrmT10A substrate tRNA^Gln^ lacking m^1^G9 to become susceptible to tRNA fragmentation (Igoillo-Esteve et al. 2013; Cosentino et al. 2018). The accumulation of 5’-tRFs derived from this tRNA, while 3’-tRF levels remained largely the same in patient and control cells, suggests a role for these fragments in the human disease. Nonetheless, while the presence of the m^1^G9 modification is clearly implicated in human health, the molecular basis for the specific disorders experienced by humans with disease-associated Trm10 mutations and how human tRNAs are impacted by the lack of m^1^G9 are not fully understood. A better understanding of this enzyme can help in identifying the molecular basis for the human health impacts of Trm10 deficiency, and may also reveal principles that are applicable to other tRNA modification-related diseases.

In *S. cerevisiae*, where the activity of Trm10 was first characterized, there is only one homolog of Trm10 that performs the m^1^G9 modification on 13 of the 24 G9-containing tRNA species whose modification status is known (three G9-containing tRNA species remain uncharacterized in terms of modifications to date) (Jackman et al. 2003; Swinehart et al. 2013). *TRM10* in *S. cerevisiae* is not an essential gene, but deletion of *trm10* causes growth hypersensitivity to low concentrations of the antitumor drug 5-fluorouracil (5FU), which was revealed by a screen of the yeast deletion collection for strains that exhibit hypersensitivity to growth in the presence of 5FU (Gustavsson and Ronne 2008). Interestingly, in the genome-wide survey, the most 5FU-sensitive strains were a group of rRNA and tRNA modifying enzyme gene deletions, adding to evidence suggesting that the effect of 5FU on structure and/or function of non-coding RNAs is associated with the toxicity of this heavily used drug for treatment of solid cancer tumors (Hoskins and Butler 2008; Bash-Imam et al. 2017; Ge et al. 2017). 5FU is known to be directly incorporated into RNA, but also inhibits pseudouridine modification, which is normally abundant and functionally important in tRNA and other RNA molecules (Hoskins and Butler 2008; Borchardt et al. 2020). Thus, the loss of pseudouridine is likely to explain some of the sensitivity of RNA-associated genes to growth in the presence of the drug. However, only a subset of tRNA modification enzymes were identified to have the 5FU-hypersensitive phenotype when deleted, suggesting that the hypersensitivity is not due to a global effect on the tRNA pool, but rather that there may be specifically targeted tRNA species or structures that are uniquely sensitive to the presence of 5FU and lack of certain modifications (Gustavsson and Ronne 2008). Among the 5FU hypersensitive deletion strains identified, the *trm10Δ* strain exhibited the most severe growth defect in the presence of 5FU, suggesting that the m^1^G9 modification must be important for the ability of one or more tRNAs to withstand the effects of 5FU toxicity, but the impact of m^1^G9 on any of its substrate tRNAs in *S. cerevisiae* has not been demonstrated to date. Here, we sought to define the biological importance of the highly conserved m^1^G9 modification by taking advantage of the 5FU hypersensitive phenotype in *S. cerevisiae*.

In this study we demonstrate that the hypersensitivity of the *trm10Δ* strain to 5FU is due to severely depleted levels of a single Trm10 substrate, tRNA^Trp^. Moreover, levels of mature tRNA^Trp^ are significantly decreased in the *trm10Δ* strain even in the absence of 5FU, despite no obvious growth defect of the *trm10Δ* strain. We also observed that levels of pre-tRNA^Trp^ increase significantly upon deletion of *trm10* compared to wild-type cells. We identified that *trm10Δ* hypersensitivity to 5FU and decreased levels of hypomodified mature tRNA^Trp^ are rescued by deletion of *met22*, an enzyme that has been previously associated with at least two tRNA quality control pathways (RTD and MPD). The double mutant *trm10Δ met22Δ* strain grows normally in the presence of 5FU and its levels of mature tRNA^Trp^ are rescued to wild-type abundance. We sought to define the specific *MET22*-dependent pathway that was being used to degrade tRNA^Trp^ lacking the m^1^G9 modification by creating double mutants of *trm10Δ* with several known exonucleases, including *XRN1*, *RAT1*, and *RRP6* yet surprisingly none of these rescued the 5FU growth hypersensitive phenotype. Taken together, our results suggest that unknown nuclease(s) associated with *met22* sense tRNA^Trp^ lacking the m^1^G9 modification in *S. cerevisiae*, providing insight into how the cell recognizes and accounts for aberrant tRNA, while providing another example of a tRNA modifying enzyme that modifies numerous substrates in the cell but is only important for the function of one tRNA.

## RESULTS

### Overexpression of tRNA^Trp^ rescues growth hypersensitivity of *trm10Δ* to 5-fluorouracil

We took advantage of the strong 5FU hypersensitive phenotype of *trm10Δ* strains to study the significance of the m^1^G9 modification *in vivo* in *S. cerevisiae*. We employed an approach that has been used to study other (mainly temperature-sensitive) phenotypes associated with loss of tRNA modifications, and individually overexpressed 39 different elongator tRNAs from high copy number (2μ) plasmids in the *trm10Δ* strain to determine whether they could complement the 5FU hypersensitive phenotype (Chernyakov et al. 2008; Han et al. 2015). These tRNAs include all 12 elongator tRNA substrates of Trm10, as well as 27 other A9 and G9-containing non-substrate tRNAs, with the idea that if the abundance of a specific tRNA is impacted by the loss of m^1^G9 modification, adding more of the affected tRNA (despite its lack of modification) back to the cell will rescue growth. Of the 12 elongator tRNAs that are substrates for Trm10, only overexpression of tRNA^Trp^ was able to successfully rescue growth of the *trm10Δ* strain in the presence of increasing concentrations of 5FU, including at the highest tested concentration (5 μg/ml) where there is no detectable growth of the *trm10Δ* cells (**Figure 1A**). Four tRNAs (tRNA^Asn(GUU)^, tRNA^Cys(GCA)^, tRNA^Thr(AGU)^ and tRNA^Val(UAC)^) are not substrates for Trm10 in wild-type yeast, but are capable of being modified by purified Trm10 enzyme *in vitro* (Swinehart et al. 2013). However, none of these four tRNAs rescued the 5FU phenotype (**Figure 1A**). As expected, overexpression of any of the G9-containing tRNAs that are unmodified by Trm10 had no detectable effect on the ability of the *trm10Δ* strain to grow in the presence of 5FU (**Figure 1A**). As a control, a selected group of substrate and non-substrate tRNAs were overexpressed in the wild-type *TRM10* background (**Figure 1B**). Here, none of the overexpressed tRNAs (including tRNA^Trp^) conferred any detectable growth advantage at the highest concentrations of 5FU tested. These results indicate that there is no general ability of individual tRNAs (including tRNA^Trp^) to confer 5FU resistance when tRNAs contain the m^1^G9 modification. Likewise, no increased resistance to 5FU was observed upon overexpression of any A9-containing tRNAs, which are also not substrates for Trm10 modification in *S. cerevisiae* (**Figure 1C**). Even though overexpressed tRNA^Trp^ remains unmodified in the *trm10Δ* background, we hypothesize that increased abundance of this tRNA is sufficient to rescue growth in the presence of 5FU, leading us to further examine whether the *trm10Δ* hypersensitivity to 5FU is caused by decreased tRNA^Trp^ levels that are insufficient for the cell.

**Figure 1.**
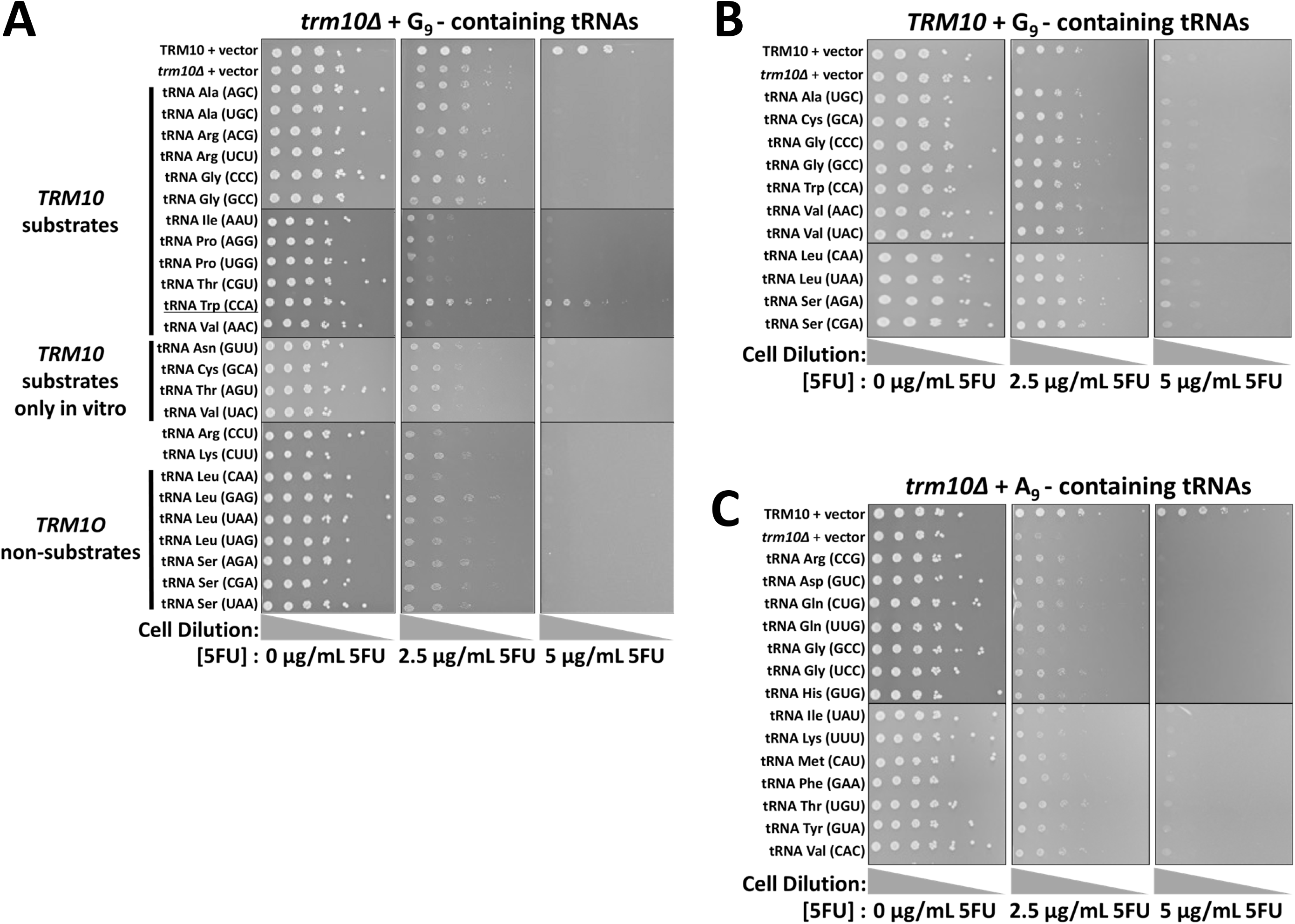
Overexpression of Trm10 substrate tRNA^Trp^ rescues the *trm10Δ* growth defect in 5FU. Strains were grown overnight in SD-Leu media, serially 4-fold diluted from a starting OD_600_ of 1, plated on SD-Leu plates with indicated concentrations of 5FU, and incubated for 4 days at 30°C. **(A)** G_9_-containing tRNAs overexpressed in the *trm10Δ* strain. Whether the tRNA is modified *in vivo* or only by purified Trm10 *in vitro* is indicated on the left of each figure. The modification status of tRNA^Arg(CCU)^ has not been determined in *S. cerevisiae*. *S. cerevisiae* tRNA^Lys(CUU)^ contains m^2^G9, and can be modified by Trm10 upon overexpression of the enzyme *in vivo*, but has not been tested further as a Trm10 substrate *in vitro*. **(B)** G_9_-containing tRNAs overexpressed in *TRM10* cells. **(C)** A_9_-containing tRNAs overexpressed in *trm10Δ* cells.

### Levels of mature tRNA^Trp^ are significantly depleted in *trm10Δ* strains

To determine whether the inability of the *trm10Δ* strain to grow in the presence of 5FU is correlated with the abundance of tRNA^Trp^, we performed northern analysis using a probe targeting mature tRNA via its anticodon stem and D-loop sequences, which lack other modifications that would predictably block primer binding. Interestingly, we observed significantly lower levels of mature tRNA^Trp^ in the *trm10Δ* strain compared to the isogenic *TRM10* control, even in the absence of 5FU (**Figure 2A** compare lane 7 with lane 3, quantified in **Figure 2B**). Therefore, even though there is not a detectable growth phenotype of the *trm10Δ* strain in the absence of 5FU, levels of mature tRNA^Trp^ are already significantly decreased compared to the wild-type strain. Consistent with the 5FU hypersensitivity of the *trm10Δ* strain, the added stress of growth in 5FU further depleted levels of mature tRNA^Trp^ in the *trm10Δ* strain to extremely low levels (<5% of the amount in the wild-type strain) (**Figure 2B**). Presumably, the abundance of tRNA^Trp^ in *trm10Δ* cells grown in 5FU is not sufficient to sustain translation and therefore viability. In agreement with the rescued growth hypersensitivity to 5FU upon tRNA^Trp^ overexpression, mature tRNA^Trp^ levels increased in each of the tRNA^Trp^ overexpressing strains. Although the abundance of tRNA^Trp^ does not fully recover to wild-type levels in the *trm10Δ* background under any condition, the levels measured in the *trm10Δ* strain with tRNA^Trp^ overexpressed +5FU are similar to those observed in the *trm10Δ* strain -5FU where there is no growth phenotype (**Figure 2B**). The inability of tRNA^Trp^ to reach wild-type levels in these strains is likely due to the fact that the tRNA remains unmodified, and therefore is still likely subject to the action of the quality control pathways that act to remove the incorrectly modified tRNA from the cells.

**Figure 2.**
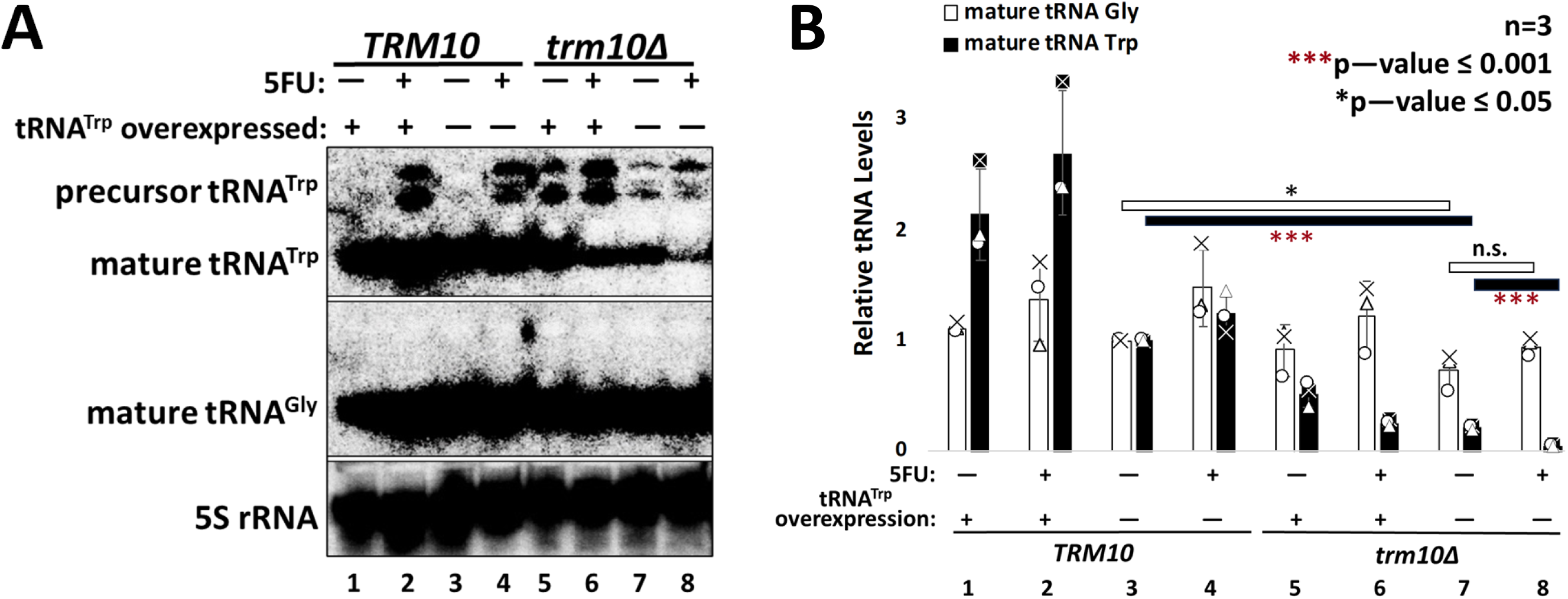
Levels of tRNA^Trp^ decrease upon deletion of *trm10* and are almost entirely depleted upon growth of this strain in 5FU. **(A)** Northern analysis of RNA derived from the indicated strains grown with and without 5FU. Mature tRNA^Trp^, mature tRNA^Gly(GCC)^, and 5S rRNA levels were determined using 5’ radiolabeled probes (**Table S2**). **(B)** Quantification of relative tRNA levels from strains shown in (A). Relative tRNA levels were calculated by comparing the observed intensity for each tRNA to the normalized abundance observed in the *TRM10* strain (lane 3) set to 1. Triplicate data was plotted with each data point shown; error bars denote standard deviation. Two sample t-tests assuming equal variances were performed, and one-tailed P-values shown with red asterisks under black bars for comparison of tRNA^Trp^ levels or a black asterisk or n.s. (no significant difference) over white bars for comparison of tRNA^Gly(GCC)^ levels. Specific p-values from other t-test comparisons are shown in **Table S1**.

As a control, we performed northern blots targeting another Trm10 substrate, tRNA^Gly(GCC)^ whose overexpression in the *trm10Δ* strain does not rescue 5FU growth hypersensitivity (**Figure 1A**). As expected, quantification of the northern data did not reveal any dramatic changes in the abundance of tRNA^Gly(GCC)^ in any of the tested strains (**Figure 2B**). We reproducibly observed a small, but statistically significant (p ≤ 0.05), decrease in abundance of tRNA^Gly^ in *trm10Δ* cells when compared to *TRM10* cells grown under the same conditions (**Figure 2B**, white bar comparing *TRM10* vs. *trm10Δ* in the absence of 5FU, and **Table S1** listing p-values for other comparisons). The addition of 5FU to the media has, if anything, a slightly positive effect on the abundance of tRNA^Gly(GCC)^, although statistical analysis indicates that none of the apparent changes rise to the level of significance observed with tRNA^Trp^. These results indicate that tRNA^Trp^ alone remains more greatly affected by loss of the m^1^G9 modification, consistent with its unique ability to restore growth in 5FU (**Figure 2B** and **Table S1**). Moreover, unsurprisingly, the overexpression of tRNA^Trp^ does not detectably affect the abundance of tRNA^Gly^ under any tested condition. Overall, the observed pattern of mature tRNA levels suggests that the presence of the m^1^G9 modification is selectively important for maintaining sufficient levels of tRNA^Trp^ under 5FU growth conditions.

### Levels of pre-tRNA^Trp^ accumulate in a Trm10- and 5FU-dependent manner

Hybridization of the tRNA^Trp^ probe that targets sequences in the mature tRNA also revealed the presence of higher migrating precursor tRNA^Trp^ (pre-tRNA^Trp^) species in several strains (**Figure 2A**). We note that the tRNA^Gly(GCC)^ oligo hybridizes to a similar region of the mature tRNA, yet did not similarly indicate the presence of pre-tRNA^Gly^ in any of the strains (**Figure 2A**). To further assess potential tRNA processing defects associated with these conditions, we probed the same RNAs with oligonucleotide probes designed to target specific pre-tRNA^Trp^ species (Chatterjee et al. 2022). Two probes targeted two of the six tRNA^Trp^ 5’-leader sequences (named GAT and GTT based on the last three nucleotides of the distinct 5’-leader sequences for each tRNA^Trp^ gene), and one probe targeted intron-containing pre-tRNA^Trp^ (**Figure 3; Table S2; Table S3**). Notably, all precursor probes included some exon sequence to facilitate sufficient hybridization to the target RNA; therefore, mature tRNA is also detected in each of these assays to varying extents (as described in more detail below).

**Figure 3.**
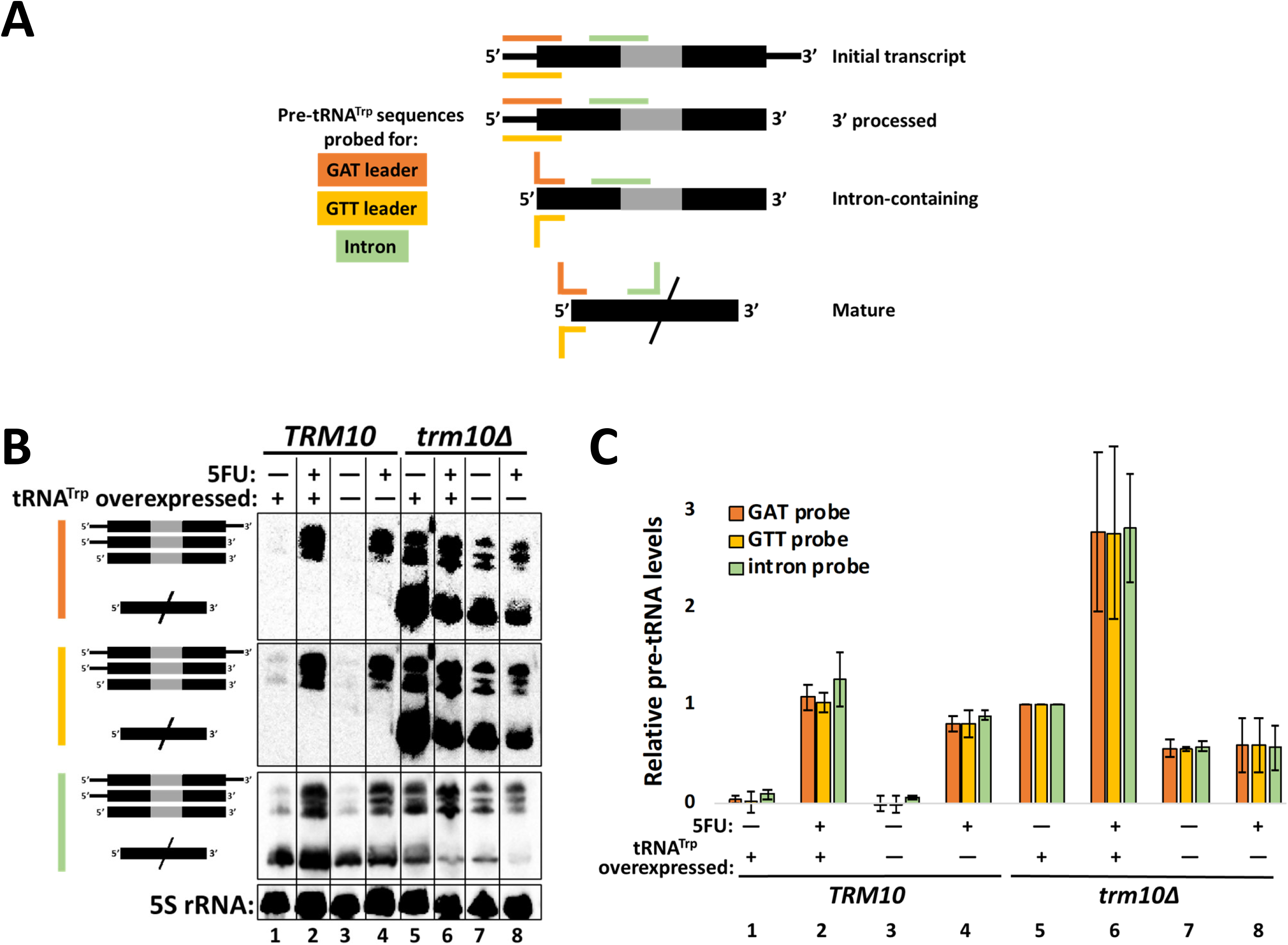
tRNA^Trp^ precursors accumulate in *trm10Δ* strains and in *TRM10* strains grown in 5FU. **(A)** Diagram indicating the three pre-tRNA^Trp^ species detected by northern probes. Orange and yellow lines indicate hybridization of 5’ leader probes (GAT and GTT, indicating the last three nucleotides of the targeted leader sequence, respectively), while green lines indicate hybridization of the intron probe (sequences listed in **Table S2**). Images are not to scale, but bent lines indicate probe sequences that lose hybridization to intron-containing and mature tRNAs. **(B)** Northern analysis of RNA dervied from the indicated strains using 5’-biotinylated probes and visualized with chemiluminescence. The bar to the left of each set of results indicates the probe that was utilzied (colored as in (A)). **(C)** Quantification of relative total pre- RNA levels (for all pre-tRNA species combined) from strains shown in (B). Relative tRNA levels were calculated by comparing the observed intensity for each RNA to the normalized abundance observed in the *trm10Δ* strain plus tRNA^Trp^ overexpression (lane 5), to allow for quantifying the full range of increased and decreased pre-tRNA levels. Duplicate data was plotted, with each end of the error bars showing each data point.

tRNA^Trp^ is unique in *S. cerevisiae* in that it is the only tRNA that is processed by removal of the 3’ trailer sequence prior to removal of the 5’ leader sequence (O’Connor and Peebles 1991; Kufel and Tollervey 2003). Therefore, the three species that are observed in northerns with pre-tRNA probes are 1) the initial transcript that contains a 5’ leader, 3’ trailer, and an intron, 2) the 3’ processed (5’ leader- and intron-containing) pre-tRNA^Trp^, and finally, 3) the end-processed, but still intron-containing pre-tRNA^Trp^ that will be exported to the cytoplasm, where tRNA splicing takes place on the surface of the mitochondria (**Figure 3A**). The absence of a band for mature tRNA^Trp^ in *TRM10* cells observed with either of the 5’-leader-targeting probes is explained by the fact that these probes both overlap the G9 nucleotide (**Figure 3B**). The m^1^G9 modification in *TRM10* cells blocks base-pairing, and in the *trm10Δ* strains that lack m^1^G9 modification, mature tRNA^Trp^ is readily detected with both 5’-leader hybridizing probes (**Figure 3B**, compare the mature tRNA band in lane 3 vs. lane 7 for both leader probes). Consistent with this result, the intron probe readily detected mature tRNA^Trp^ from both *TRM10* and *trm10Δ* cells, with levels of mature tRNA^Trp^ that mirror those measured with the mature tRNA-targeting probe used in Figure 2 that does not overlap G9 (**Figure S1D**).

Pre-tRNA^Trp^ species do not accumulate in *TRM10* cells grown in the absence of 5FU, even upon tRNA^Trp^ overexpression, suggesting that accumulation of pre-tRNA^Trp^ cannot be simply attributed to overwhelming the tRNA processing machinery with excess tRNA^Trp^ (**Figure 3B**, lanes 1 and 3). However, addition of 5FU to *TRM10* cells causes significant pre-tRNA^Trp^ accumulation even though the m^1^G9 modification is present, suggesting a processing defect for tRNA^Trp^ that is likely caused by incorporation of 5FU into the tRNA, separate from the effects of Trm10. Interestingly, however, deletion of *trm10* alone also causes significant accumulation of precursor tRNA^Trp^ levels, which is observed even in the absence of 5FU (**Figure 3B**, compare lanes 3 and 7). Similar patterns of total pre-tRNA^Trp^ accumulation with distinct Trm10- and 5FU-dependence were observed upon quantification of results with all three pre-tRNA directed oligos (**Figure 3C**).

Overall, in the presence of *TRM10*, pre-tRNA^Trp^ species accumulate to similar levels in the presence of 5FU regardless of tRNA^Trp^ overexpression (**Figure 3B, 3C**, lanes 1 and 2 vs. lanes 3 and 4). However, the results in the context of *trm10* deletion are very different, where overexpression of tRNA^Trp^ leads to a large 5FU-dependent increase in pre-tRNA^Trp^ (**Figure 3B**, lane 5 vs. 6), but pre-tRNA^Trp^ accumulation remains unchanged by the addition of 5FU to the t*rm10Δ* strain alone (**Figure 3B**, lane 7 vs. 8). These data indicate that the loss of m^1^G9 causes an accumulation of tRNA^Trp^ precursors that is separate from (and additive with) the effects of 5FU.

To test whether the 5FU or *trm10Δ* effects could be attributed to a specific processing defect, the northern intensities of each of the three individual pre-tRNA species were quantified separately (**Figure S1A, B,** and **C**). All three pre-tRNA targeting probes showed the same pattern of pre-tRNA abundance changes on both 5’-leader containing pre-tRNA species: the initial precursor and the 3’-processed (5’-extended) pre-tRNA, reflecting the fact that these pre-tRNAs can completely hybridize to all three DNA probes (**Figure 3A**). Neither deletion of *trm10* or addition of 5FU significantly perturbs the 3’-end processing step for pre-tRNA^Trp^, since we did not observe higher relative levels of initial transcript (**Figure S1A**) at the expense of 3’-processed species (**Figure S1B**) when comparing levels in any of the same strains. However, the amounts of end-matured intron-containing pre-tRNA reveal some intriguing differences (**Figure S1C**). To look more quantitatively at these levels, we compared relative levels measured with the intron-containing probe, since end-matured intron-containing pre-tRNA^Trp^ completely hybridizes to the intron probe, but not the two leader probes (**Figure 3A**). Interestingly, both 5’-leader containing species (**Figure S1A and S1B**) accumulate to substantially higher levels than intron-containing pre-tRNA (**Figure S1C**) in *TRM10* strains +5FU (**Table S4**). The 5FU-dependent increase in 5’-extended pre-tRNA is ∼18-37-fold in these strains, whereas the intron containing pre-tRNA only increases ∼7-fold in *TRM10* strains +5FU (**Table S4)**. Thus, 5FU appears to preferentially impair 5’-leader removal from pre-tRNA^Trp^, leading to relatively more accumulation of 5’-extended precursors as a fraction of the total pre-tRNA species in these cells. Separately, comparison of individual pre-tRNA species between *TRM10* and *trm10Δ* strains reveals a more modest accumulation of 5’-extended transcripts compared to intron-containing pre-tRNA (**Figure S1, Table S4**), suggesting that the negative effect on pre-tRNA^Trp^ processing observed in *trm10Δ* strains cannot be as strongly attributed to a specific defect in a particular step of the pre-tRNA processing pathway.

### Deletion of *met22* rescues growth hypersensitivity of *trm10Δ* strains to 5FU

To identify the pathway by which tRNA^Trp^ levels are reduced upon loss of m^1^G9 modification, we created double deletion strains of *trm10Δ* and genes coding for enzymes associated with several known tRNA quality control pathways in *S. cerevisiae.* The rationale for these experiments is that inactivation of a tRNA quality control pathway that degrades hypomodified tRNA^Trp^ would stabilize the tRNA and rescue the 5FU growth defect, as has been observed for other targets of tRNA surveillance. We used homologous recombination to replace each gene. The *trm10Δ* strain available from the yeast genome deletion collection has a kanamycin-resistant (kanMX) cassette replacing the *TRM10* coding sequence, enabling us to introduce alleles of *met22*, *xrn1* and *rat1* that had been constructed with other drug-resistant cassettes (see **Table 1** for full strain genotypes) (Chernyakov et al. 2008))(Chernyakov et al. 2008). However, we also constructed a *trm10Δ::natMX* strain, and demonstrated that it exhibits the same 5FU hypersensitivity of the *trm10Δ::kanMX* deletion strain (**Figure 4A**), to enable the use of kanMX deletion alleles to target other quality control genes that were already available from the yeast deletion collection.

**Figure 4.**
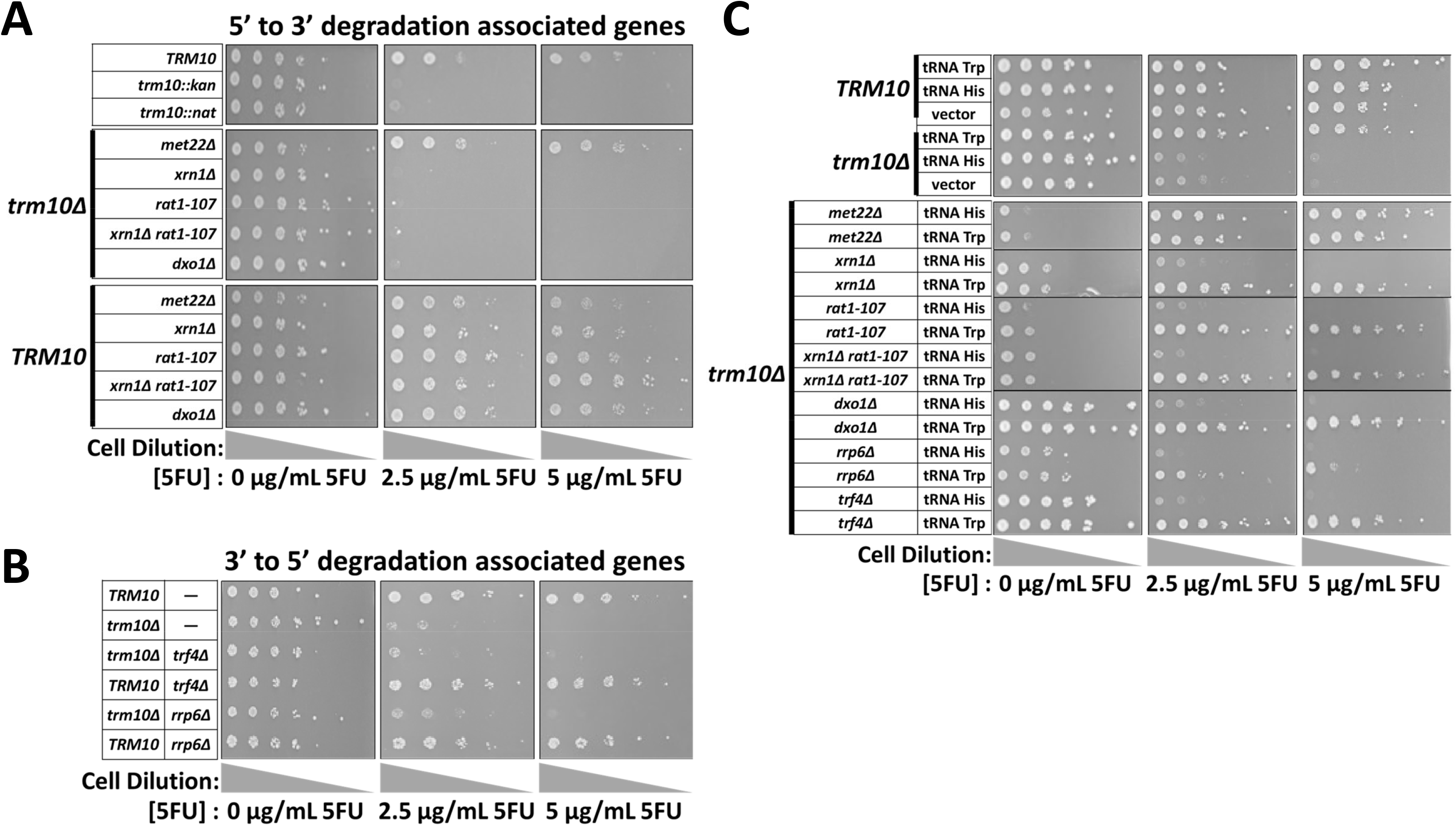
Deletion of *met22* in *trm10Δ* strain rescues growth sensitivity to 5FU. Strains were grown overnight in SC or SD-Leu media (if containing a tRNA overexpression plasmid), serially 4-fold diluted from a starting OD_600_ of 1, plated on plates of corresponding media with indicated concentrations of 5FU, and incubated for 4 days at 30°C. **(A)** Double and triple mutants with *trm10Δ* and other possible genes that have been implicated in 5’-3’ tRNA decay were tested for ability to revert 5FU hypersensitivity; control *TRM10* strains are shown with the same mutations, as indicated. Note that the *trm10Δ* 5FU hypersensitive phenotype is observed with both kanR and natR gene replacements (see control strains at top of panel). **(B)** Double mutants of *trm10Δ* with genes associated with 3’-5’ tRNA decay, and control *TRM10* strains with the same mutations, as indicated. **(C)** Overexpression of tRNA^Trp^ or tRNA^His^ from plasmids was performed in the indicated *TRM10* or *trm10Δ* strain backgrounds. The results with the vector control strain in *TRM10* and *trm10Δ* cells are shown in the top panel.

**Table 1.**
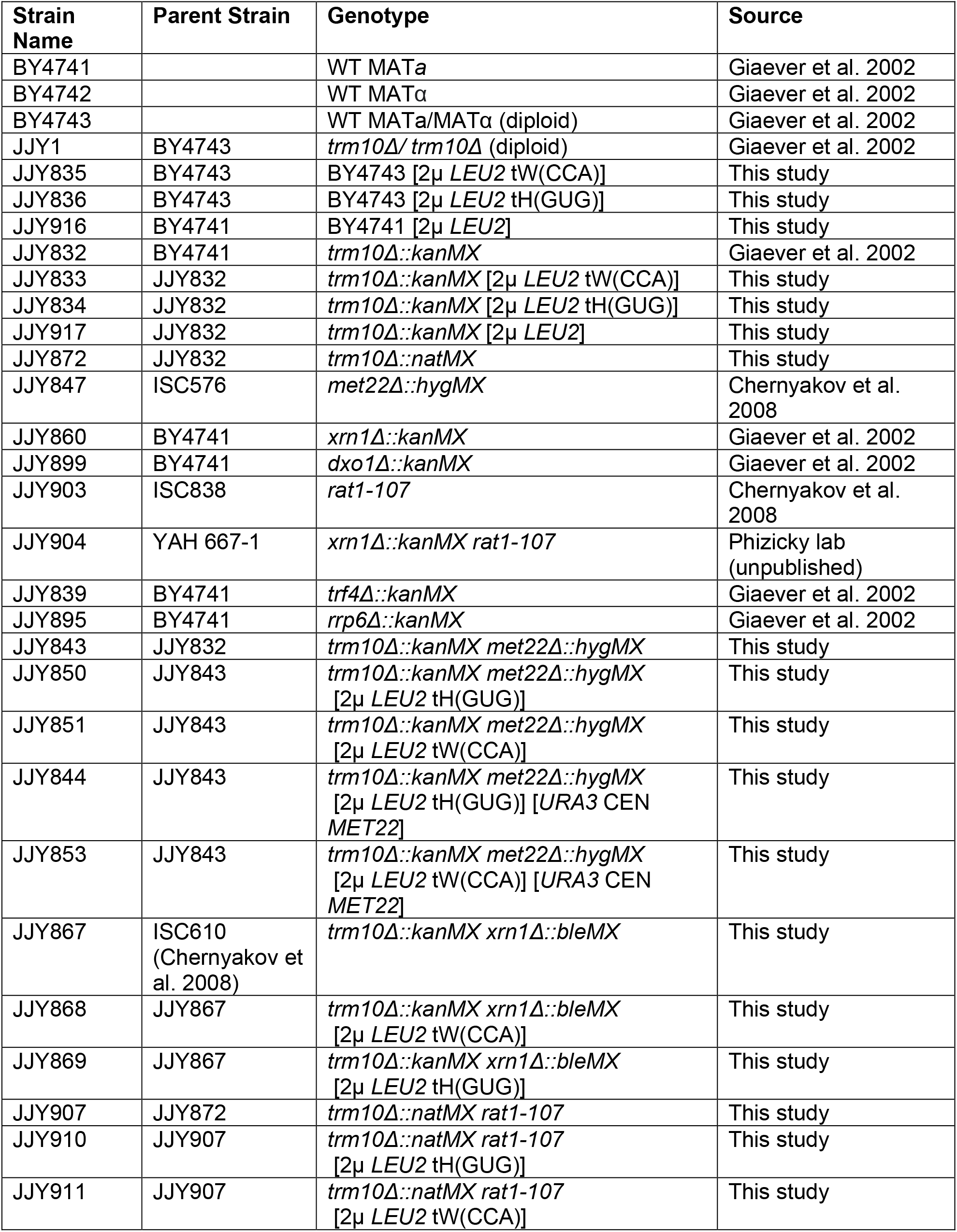
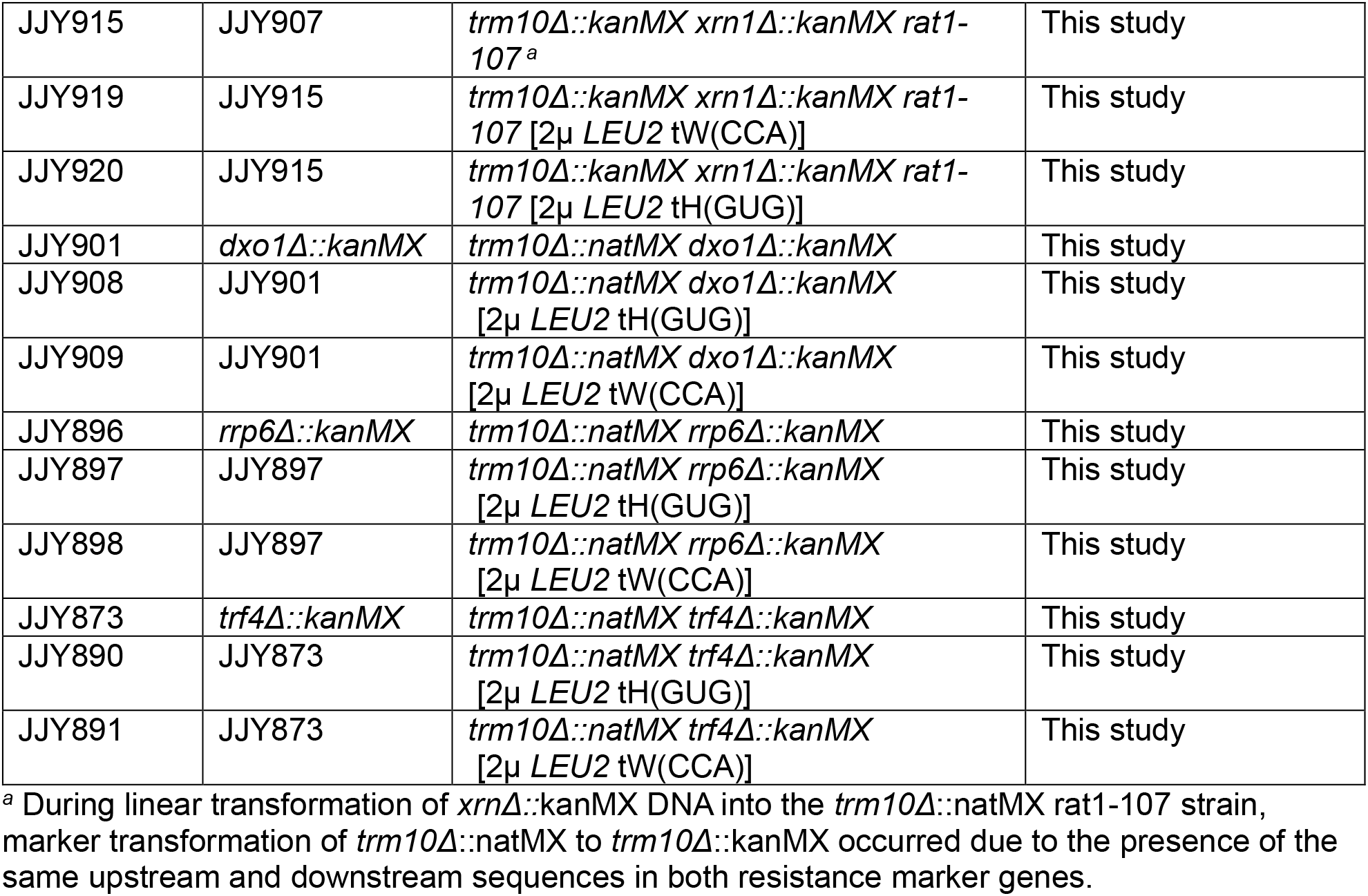
*S. cerevisiae* strains used in this study.

Met22 is an enzyme of methionine biosynthesis that, when deleted, causes accumulation of its substrate, adenosine 3’,5’ bis-phosphate (pAp). This metabolite is a potent inhibitor of the 5’ to 3’ exonucleases Xrn1 and Rat1, which have both been shown to degrade aberrant tRNA in the context of the RTD quality control pathway (Murguia et al. 1996; Dichtl et al. 1997). Targets of RTD are stabilized in the presence of *met22Δ* because accumulated pAp inhibits these 5’ to 3’ exonucleases (Alexandrov et al. 2006; Chernyakov et al. 2008). Double deletion of *trm10* and *met22* rescued growth in the presence of 5FU (**Figure 4A**). The observed growth rescue by *met22Δ* could have been consistent with RTD-mediated degradation of mature tRNA^Trp^ that lacks m^1^G9, which is exacerbated in the presence of 5FU. To test this, we created double mutant strains with *trm10Δ* and either or both of the RTD-associated 5’-3’ exonucleases, *XRN1* and *RAT1*. *XRN1* is a non-essential gene in *S. cerevisiae*, allowing us to create *trm10Δ xrn1Δ* double deletion strains, but *RAT1* is essential, so we introduced a *rat1-107* allele used previously to demonstrate the role of this enzyme in degradation of specific hypomodified tRNA by RTD (Chernyakov et al. 2008). Interestingly, unlike for other RTD quality control mechanisms discovered to date, neither deletion alone was able to reverse the *trm10Δ* phenotype, as both *trm10Δ xrn1Δ* and *trm10Δ rat1-107* strains exhibited the same hypersensitivity to 5FU as the single *trm10Δ* mutant (**Figure 4A**). To test possible redundancy of Xrn1/Rat1 in tRNA^Trp^ surveillance, we created the triple *trm10Δ xrn1Δ rat1-107* strain, but again, observed no restoration of 5FU-resistant growth. This led us to reason that the effects of *met22Δ* are occurring through another, yet unidentified, Met22-dependent mechanism for tRNA surveillance and removal that does not require either Xrn1 or Rat1 5’-3’ exonuclease activity. Another *S. cerevisiae* 5’-3’ exonuclease (*Dxo1*) has not been associated with *met22Δ* or observed to act on tRNA. Nonetheless, because of its mechanistic similarity to the RTD exonucleases, we also constructed the *trm10Δ dxo1Δ* strain (Chang et al. 2012; Yun et al. 2018). Again, we observed no change in 5FU hypersensitivity, ruling out this enzyme as the Met22-dependent enzyme that is acting on tRNA^Trp^ (**Figure 4A**). We note that Met22-dependent degradation of intron-containing pre-tRNA by an as of yet unidentified nuclease has been identified in *S. cerevisiae*, but since that pathway senses pre-tRNA, a distinct pathway must participate in mature tRNA^Trp^ decay described here (Tasak and Phizicky 2022). The TRAMP complex also degrades hypomodified tRNA^iMet^ in *S. cerevisiae*, but neither deletion of *trf4* (an essential component of the TRAMP complex) or the *rrp6* subunit of the nuclear exosome that degrades the tRNA after polyadenylation by the TRAMP complex was able to rescue the 5FU growth defect of the *trm10Δ* strain, ruling out a role for the TRAMP complex in tRNA^Trp^ quality control (**Figure 4B**) (Kadaba et al. 2004; Callahan and Butler 2010). The lack of even partial growth rescue in the *trm10Δ trf4Δ* and *trm10Δ rrp6Δ* strains also rules out a mechanism involving both RTD and TRAMP, as was recently observed with tRNA^iMet^ (Tasak and Phizicky 2022).

To rule out the possibility that there is a new cause for 5FU hypersensitivity in the context of the double and triple deletion strains tested in **Figure 4**, we overexpressed tRNA^Trp^ and a control tRNA^His^ (which is not a Trm10 substrate and does not affect 5FU sensitivity) in each double and triple deletion strain (**Figure 4C**). In all cases, transformation of the plasmid directing overexpression of tRNA^Trp^, but not tRNA^His^, was able to restore wild-type levels of growth in the presence of 5FU in the double and triple mutant strains, as is observed for *trm10Δ* alone. These results indicate that insufficiency of tRNA^Trp^ remains the cause of 5FU hypersensitivity that persists in each strain, and not that another tRNA becomes limiting in the absence of the other tRNA quality control enzymes. Therefore, we conclude that mature tRNA^Trp^ lacking the m^1^G9 modification is recognized in a novel manner by the cell for degradation. Growth hypersensitivity of *trm10Δ* to 5FU is rescued by deletion of *met22*, yet not by inactivation of any of the known exonuclease-dependent pathways that have been associated with Met22 to date.

### Growth rescue in the *met22Δ trm10Δ* strain correlates with restored abundance of mature tRNA^Trp^

Although decreased abundance of tRNA^Trp^ appears to remain the barrier to viability in the double mutant strains in the presence of 5FU (**Figure 4C**), the lack of an effect of Xrn1, Rat1 or Trf4 deletion raised questions about whether there is another previously undescribed Met22-associated mechanism that does not involve inhibiting tRNA decay due to pAp accumulation. Thus, we quantified mature tRNA^Trp^ and tRNA^Gly^ levels in RNA isolated from the *trm10Δ met22Δ* strain under each of the previously tested growth conditions, with and without *MET22* expressed in the strain (**Figure 5**).

**Figure 5.**
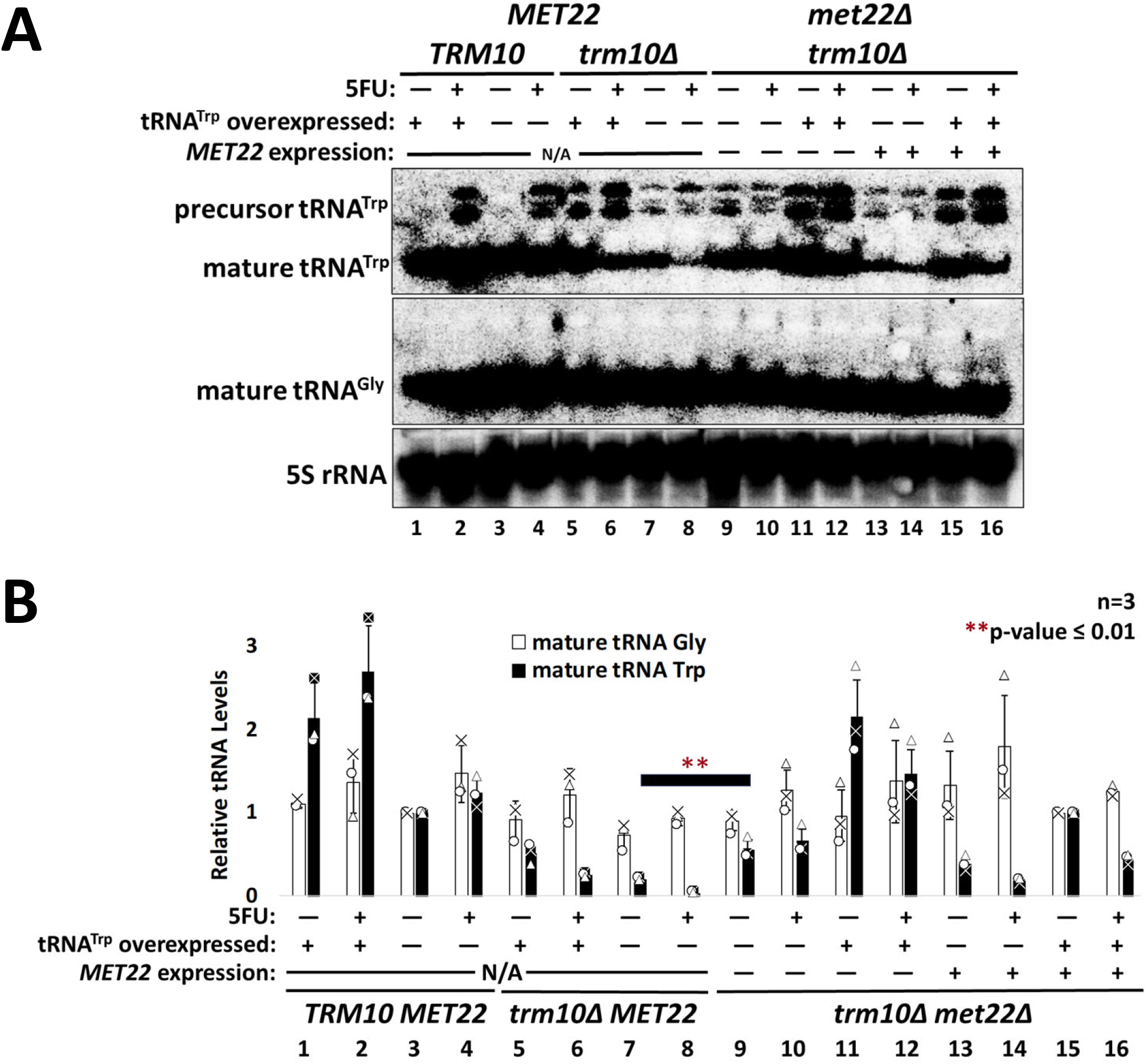
Mature tRNA^Trp^ levels are rescued upon *met22* deletion in *trm10Δ* strain. **(A)** Northern analysis of RNA derived from the indicated strains probed for mature tRNA^Trp^, mature tRNA^Gly^ and 5S rRNA, as indicated. **(B)** Quantification of relative mature tRNA levels from strains shown in (A). Normalized RNA levels (to 5S rRNA) were calculated for each strain, and for lanes 1-8, lane 3 was set to 1 for determining relative tRNA levels, and for lanes 9-16, lane 15 was set to 1 for determining relative tRNA levels, in order to better visualize the positive and negative changes associated with each set of strains on the same axis. Triplicate data was plotted, with individual data points for tRNA^Gly(GCC)^ and tRNA^Trp^ shown. Two tailed t-tests assuming equal variance were performed, and one-tailed P-value is represented by asterisks comparing tRNA^Trp^ levels in the *trm10Δ* and the *trm10Δmet22Δ* strain. P-values for comparisons not shown here are listed in **Table S1**.

In *trm10Δ met22Δ* strains, levels of mature tRNA^Trp^ increased compared to levels observed in the corresponding *trm10Δ* single mutant strains (**Figure 5B**, compare mature tRNA^Trp^ levels in lanes 9 and 10 vs. in lanes 7 and 8). This remains consistent with the known mechanism for Met22 that involves inhibition of 5’-3’ exonucleases that degrade mature hypomodified tRNA, despite the fact that neither of its known RTD-associated enzymes appear to be involved in quality control of tRNA^Trp^. Interestingly, although expression of *MET22* from this *CEN* plasmid did not restore the 5FU hypersensitive growth phenotype to the *trm10Δ met22Δ* strain (data not shown), the abundance of tRNA^Trp^ was detectably decreased by restoration of active Met22 in the complemented strain (**Figure 5**, for example, lanes 13 vs. 9 and 14 vs. 10). Plasmid-borne Met22 thus partially complements the molecular phenotype (decay of tRNA^Trp^) but appears not to be expressed to sufficient levels to fully counteract the stabilizing effect of *met22Δ* on tRNA^Trp^ abundance, and thus enable reversion to 5FU sensitive growth of the *trm10Δ* strain.

Despite the ability of *met22Δ* to rescue growth and increase mature tRNA^Trp^ levels in the *trm10Δ* background, pre-tRNA^Trp^ species are still detected in the *trm10Δ met22Δ* cells (**Figure 5A**, lanes 9-12). We quantified pre-tRNA^Trp^ accumulation in the *trm10Δ met22Δ* cells by directly probing for pre-tRNA species using the same oligonucleotide probes as in Figure 2 (**Figure S2**). These data confirmed that although the expression of *MET22* in the *trm10Δ met22Δ* background resulted in significant destabilization of the mature tRNA^Trp^ (**Figure S2A**, compare lanes 1 vs. 5, 2 vs. 6, 3 vs. 7, or 4 vs. 8), the relative abundance of pre-tRNA species was not significantly affected in the same strains (**Figure S2**). The much more substantial impact of *met22Δ* on mature tRNA^Trp^ levels rather than on pre-tRNA accumulation suggests that 5FU hypersensitivity of the *trm10Δ* strain is mostly driven by degradation of the mature tRNA rather than by the defect in pre-tRNA processing.

## DISCUSSION

Here we used genetic and biochemical approaches to demonstrate that the previously observed 5FU hypersensitive phenotype associated with deletion of the tRNA m^1^G9 methyltransferase Trm10 *in S. cerevisiae* is uniquely due to a destabilizing effect on tRNA^Trp^, but not on other Trm10 substrate tRNAs (**Figures 1** and **2**). Moreover, the negative effect on tRNA^Trp^ is also detected in the absence of m^1^G9 modification alone, and is further exacerbated when *trm10Δ* cells are grown in the presence of 5FU (**Figures 1** and **2**). We also discovered that growth in the presence of 5FU causes a processing defect in tRNA^Trp^ that results in accumulation of partially processed tRNA^Trp^ species (**Figure 3**). Separately, *trm10Δ* strains also accumulate tRNA^Trp^ precursors even in the absence of 5FU (**Figure 3**). These data suggest that pre-tRNA^Trp^ processing steps are particularly sensitive to perturbations in the physical attributes of this tRNA. We showed that deletion of *met22* rescued the 5FU hypersensitive growth phenotype of the *trm10Δ* strain due to increased levels of mature tRNA^Trp^ in the *met22Δ trm10Δ* strain that are presumably sufficient to maintain translation even with the hypomodified tRNA (**Figure 4** and **Figure 5**). However, inactivation of other players in the known tRNA decay pathways in *S. cerevisiae* did not similarly rescue the *trm10Δ* 5FU growth defect (**Figure 4**), indicating that Met22 acts through some other yet unknown tRNA decay pathway to remove inappropriately modified tRNA^Trp^ from the cells. The fact that Trm10 modifies 13 different cytosolic tRNAs in *S. cerevisiae*, but that the biological importance of the m^1^G9 modification seems to only be significant for one of its substrates (tRNA^Trp^) provides yet another example of a tRNA modification that does not impact all the tRNAs that contain it equally (Phizicky and Alfonzo 2010; Howell and Jackman 2019). The evolutionary origins and molecular basis for the activity of Trm10 on multiple *S. cerevisiae* tRNAs remains to be fully understood.

To date, decay of mature tRNAs due to RTD has been demonstrated for three *S. cerevisiae* tRNAs, tRNA^Ser(CGA)^ and tRNA^Ser(UGA)^ and tRNA^Val(AAC)^ (Alexandrov et al. 2006; Chernyakov et al. 2008; Whipple et al. 2011; Dewe et al. 2012). Hypomodification due to genetic disruption of at least five other tRNA modification enzymes, including Trm8 (m^7^G46), Trm1 (m^2,2^G26), Trm4 (m^5^C), Tan1 (ac^4^C12) or Trm44 (Um44) in different combinations has been implicated in these tRNA quality control events, each of which impacts only one or two tRNAs from among all possible substrates for each enzyme. Interestingly, even though the abundance of some of these RTD substrates is detectably impacted upon mutation of the modification enzyme(s) alone, as we also observed for tRNA^Trp^ (**Figure 2**), growth defects are not observed until addition of another stressor to the cells, such as growth at high temperature (Dewe et al. 2012). The example we provide here is the first demonstration that growth in the presence of 5FU can similarly destabilize a tRNA that is already impacted by loss of a modification from one of its substrates. The fact that not all substrates of each modifying enzyme are similarly affected by loss of modifications underscores the challenges associated with understanding the function of individual tRNA modifications and their enzymes. These results also help to rationalize the long-standing conundrum surrounding the general lack of observed growth defects associated with genetic mutants in many of these highly conserved tRNA modification enzymes in *S. cerevisiae*, including Trm10, despite the demonstrated impact of Trm10 deficiency on human health.

The toxic effects of 5FU that cause its widespread use as an antitumor drug have been difficult to attribute to the impact on a specific nucleic acid, since 5FU is salvaged and becomes part of the total nucleotide pool. 5FU can become both an inhibitor of thymidylate synthase in the form of 5FdUMP and can be incorporated into biological RNA molecules in the form of 5FUTP. Examples of negative effects of 5FU on pre-mRNA splicing and translation have been revealed, but the demonstration that a large number of tRNA modification enzyme genes were the most sensitive to 5FU in the genome-wide deletion collection suggested that impacts of 5FU on tRNA function may be even more biologically significant (Gustavsson and Ronne 2008; Hoskins and Butler 2008; Bash-Imam et al. 2017; Ge et al. 2017). Here we provide the first molecular explanation for the effect of 5FU on tRNA function. Moreover, the observation that these effects (at least for the *trm10Δ* strain) occur predominantly on a single tRNA (Trp) despite the incorporation of 5FU into all tRNA species, is intriguing. One possible rationale for the negative impact of 5FU on RNA is due to the fact that the presence of 5FU inhibits formation of pseudouridine (Ψ) modification, which is generally thought to have a stabilizing impact on RNA and is one of the most abundant modifications of all RNAs, but is especially abundant in tRNA (Davis and Poulter 1991; Davis 1995; Hoskins and Butler 2008). The role of pseudouridylation in *trm10Δ* 5FU hypersensitivity has also been indicated by a previous study that identified a strong genetic interaction between the *trm10* and *pus3* genes (Han et al. 2015). Even though multiple tRNAs in the cell contain both Pus3-(Ψ38 and/or Ψ39) and Trm10-catalyzed modifications, only the overexpression of tRNA^Trp^ (naturally containing Ψ39, not Ψ38) partially rescued growth of the *trm10Δ pus3Δ* strain (Han et al. 2015). Together the results of this study and our work implicate loss of pseudouridylation due to incorporation of 5FU into tRNA^Trp^ as a major mechanism that contributes to the observed 5FU hypersensitive phenotype of the *trm10Δ* strain. The specific sensitivity of tRNA^Trp^ may be due to the below average predicted stability of its anticodon stem where Ψ39 modification occurs, although another Trm10/Pus3 substrate tRNA^Arg(UCU)^ is also among the tRNAs with a below-average stability anticodon stem and is not similarly affected by 5FU (Han et al. 2015). Additionally, tRNA^Trp^ contains Ψ at 6 different positions, which is the largest number of Ψ found in any of the 12 elongator tRNAs modified by Trm10. It is possible that inhibited pseudouridylation at other positions may also contribute to the observed instability tRNA^Trp^. It will be interesting to continue to characterize the structural impact of 5FU incorporation into tRNA^Trp^ and other tRNAs as the basis for the observed hypersensitive phenotypes.

The observation that *met22Δ* stabilizes tRNA^Trp^ levels implies that a similar mechanism to that observed for other Met22-dependent tRNA quality control pathways, in which accumulation of the Met22 substrate pAp inhibits exonucleases, such as Rat1 and Xrn1, that could have been acting on tRNA^Trp^. However, the identity of the nuclease(s) that act to degrade hypomodified tRNA^Trp^ remains unknown since any of the mutants of nuclease pathways that are known to participate in tRNA decay (*rat1*, *xrn1*, *trf4* or *rrp6*) do not rescue the *trm10Δ* growth defect (**Figure 4**). For other tRNAs degraded by RTD, mutation of either *xrn1* or *rat1* is sufficient to restore levels of the affected tRNA. However, the lack of 5FU growth rescue in the triple *trm10Δ xrn1Δ rat1-107* strain (**Figure 4**) confirms that quality control of tRNA^Trp^ is not just a new variant of RTD in which the 5’-3’ exonucleases are acting redundantly to degrade tRNA^Trp^. Therefore, there must be some other Met22-responsive change in tRNA decay that explains the growth rescue in the *trm10Δ met22Δ* strain.

Met22 has also been implicated in surveillance of pre-tRNA containing mutations that affect the structure of the intron-exon junction of any of several pre-tRNAs through Met22-dependent pre-tRNA decay (MPD), leading to accumulation of end matured, intron-containing pre-tRNA (Payea et al. 2015). Although deletion of *trm10* or growth in 5FU also causes accumulation of pre-tRNAs **(Figure 3)**, no further accumulation of pre-tRNAs was observed in a *met22Δ*-dependent manner that would suggest *met22Δ* is similarly acting to stabilize pre- tRNA^Trp^. Thus, MPD is ruled out as a possible mechanism for the quality control pathway for mature tRNA^Trp^ associated with *trm10Δ*. Interestingly, some crossover between RTD and the nuclear TRAMP pathway was recently observed, with initiator tRNA lacking the essential m^1^A58 modification sensed for degradation by both pathways (Tasak and Phizicky 2022). In this scenario, both *met22Δ* and *trf4Δ* were required to rescue the *trm6-504* mutation, which allows growth but lowers m^1^A58 levels. This was the first instance associating 5’ to 3’ RTD and 3’ to 5’ nuclear surveillance exonucleases in targeting the same hypomodified tRNA. Our study only found a growth rescue of the *trm10Δ* strain upon growth in 5FU with pairwise deletion of *met22* and did not detect any similar synergy between its associated pathways (**Figure 4**).

Two distinct mechanisms impact pre-tRNA^Trp^ processing, with accumulation of pre- tRNA^Trp^ observed either only in the presence of 5FU (even in *TRM10* strains) or in *trm10Δ* strains (even in the absence of 5FU). Interestingly, in *TRM10* cells grown with 5FU, although all three pre-tRNA^Trp^ precursors accumulate to some degree, accumulation of the 5’-leader containing tRNA is greater than that of end-processed intron containing tRNA (**Figure 3, S1 and Table S4**). This result suggests that there is a more substantial impact of 5FU on 5’-end processing of pre-tRNA^Trp^ than on removal of the intron. Interestingly, tRNA^Trp^ genes in *S. cerevisiae* are the only set of pre-tRNA in which all of the 6 different 5’-leader sequences contains a U at the -1 position (Chan and Lowe 2009) (**Table S3**). When cells are grown under 5FU conditions, it is likely that 5FU is incorporated into these pre-tRNA transcripts resulting in the presence of a 5FU at the -1 position in at least some tRNAs. Based on the unique processing of tRNA^Trp^ in yeast, which unlike all other pre-tRNAs already has a slower 5’-leader removal than 3’-trailer removal, it seems likely that the incorporation of 5FU at the position immediately next to Ribonuclease P-mediated 5’-end cleavage has a disproportionately negative impact on the efficiency of 5’-end leader removal causing the substantial accumulation of 5’-extended precursors. Understanding the molecular basis for the negative impact of loss of m^1^G9 on pre-tRNA^Trp^ processing is more complicated, since there is no known role for m^1^G9 in either 5’-end processing or intron splicing. It is also possible that retrograde transfer of partially processed pre-tRNA^Trp^ species also contributes to the quality control of tRNA^Trp^ by removing aberrant hypomodified pre-tRNA from the cytoplasm, as has been observed already for several other intron-containing pre-tRNA (Kramer and Hopper 2013). We note that although increased transcription of pre-tRNA^Trp^ could help to compensate for the decreased abundance of mature tRNA^Trp^, overexpression of tRNA^Trp^ in the *TRM10* wild-type strain does not cause pre-tRNA accumulation, suggesting that the defect that leads to pre-tRNA accumulation is in the processing step(s) themselves by a mechanism that remains to be determined.

## MATERIALS & METHODS

### Creation of yeast strains

Strains used in this study are listed in **Table 1**. Deletion strains generated for this study were created by amplifying the gene of interest from existing single deletion strains obtained either from the yeast genome deletion collection, or generously provided by Eric Phizicky (Giaever et al. 2002). Polymerase chain reactions (PCR) were performed using iProof polymerase (Bio-Rad) and primers designed to anneal between 150 and 400 base pairs upstream and downstream of each target gene. PCR products were purified using the NucleoSpin Gel and PCR Clean-up Kit (Machery-Nagel).

Genomic DNA for PCR amplification of target deletion genes was isolated from indicated strains as follows. Colonies of each strain were inoculated into 5mL media for overnight growth at 30°C. Cells were pelleted (5000xg for 17 minutes at room temperature), resuspended in DNA prep buffer containing 100mM NaCl, 10mM Tris HCl pH 8.0, 2% Triton X-100 and 1% SDS and then an equal volume of PCA (phenol:chloroform:isoamyl alcohol, 25:24:1) was added to the resuspended cells, followed by glass bead lysis. After vortexing at high speed for three minutes, another equal volume of TE pH 8.0 was added to the vortexed cells so buffer:PCA:1X TE is at a 1:1:1 ratio. After separation by centrifugation, the aqueous layer was transferred to a new tube, ethanol precipitation performed, and nucleic acids resuspended in 1X TE pH 8.0. 10mg/mL RNaseA (Thermo-Scientific) was added to degrade RNA in the sample. Ethanol precipitation was again performed, and the pellet resuspended in 1X TE pH 8.0 and quantified by Nanodrop prior to transformation.

Linear transformations were performed using the lithium acetate (LiOAc) method. To generate competent cells, 25 mL cultures of the target strain were grown to an OD_600_ of 1.5-2, at which point 14mL of cells were pelleted (4000 RPM for 15 minutes at room temperature) and washed two times with 0.1M LiOAc. Cells were resuspended in 140 μL 0.1M LiOAc and 32 μL salmon sperm DNA (10mg/mL Invitrogen). Purified PCR product (250-500 ng) was added to 75 μL competent cells and incubated at 30°C for 15 minutes, followed by addition of 450 μL 60% PEG 3350 1M LiOAc and further incubation at 30°C for 30 minutes. After addition of DMSO, cells were heat shocked at 42°C for 15 minutes, pelleted (4000xg for 3 minutes), and resuspended in 600 μL YPD and allowed to recover by incubation at 30°C for five hours with shaking (200 rpm). Cells were again pelleted and resuspended in 125 μL ddH_2_O and half the volume was plated on YPD plates under selective conditions for each drug resistance cassette (100 µg/mL nourseothricin sulfate (Nat), 300 µg/mL hygromycin B (Hyg), 500 µg/mL G148 (Kan), or 75 μg/mL phleomycin (Ble)). Transformants were obtained after 2-3 days of growth at 30°C and all strains were confirmed by replica plating to test for the expected drug and/or nutrient resistance, followed by PCR amplification and sequencing of the relevant gene deletion region from genomic DNA.

The *trm10Δ* strain available from the yeast deletion collection contains a kanR marker in place of the *TRM10* gene. A second *trm10Δ* strain was created using the natR marker to simplify the creation of double deletion strains using other available kanR gene replacement strains from the deletion collection. The natR gene sequence was amplified from the pAG25 vector (graciously donated by Anita Hopper’s lab) using primers designed to the common promoter/terminator regions and iProof polymerase, and transformed into the *trm10::kanR* strain as described above, with selection on YPD + Nat (100ug/mL) plates for swapping of the drug markers to create a *trm10::natR* strain.

### tRNA overexpression plasmids

A set of 2µ *LEU2* plasmids containing each of 38 different tRNA genes with their endogenous upstream and downstream genomic sequences was utilized as described in (Han et al. 2015). This collection of plasmids provided by the Phizicky lab did not contain the gene for one Trm10 substrate tRNA, and therefore, a plasmid was constructed for expression of this tRNA^Ile(AAU)^ species. This overexpression vector was created by amplifying the sequence of tRNA^Ile(AAU)^ from *S. cerevisiae* genomic DNA (YNCI0005W) with 28bp and 36bp of endogenous upstream and downstream sequence, respectively using iProof PCR. BglII and XhoI sites were introduced with the primers to enable ligation into the same AVA0577 2µ *LEU2* tRNA overexpression vector used for the rest of the tRNA overexpression collection.

### Drop tests for growth in 5FU conditions

Single colonies of each strain were inoculated in 5mL of either SD-Leu (for strains containing tRNA overexpression plasmids) or SC media and grown overnight at 30°C. Each culture was diluted to an OD_600_ of 1 in the same media, and used as the starting point for 4-fold serial dilutions of each strain. Plates containing the indicated amount of 5-fluorouracil (Sigma-Aldrich) were spotted with 2 μL of each dilution sample, and after spots were allowed to dry at room temperature, plates were incubated at 30°C. Images were taken after 3-4 days growth.

### RNA Isolation

To isolate RNA for northern analysis, cultures were grown in the absence of 5FU to late log phase and used to inoculate 1 L cultures at a starting OD_600_ of 0.01 for growth at 30°C in the presence or absence of 1 µg/mL 5FU. After 12-14 hours of growth, cultures that exhibit 5FU hypersensitivity (such as the *trm10Δ* strain) exhibit a clear growth defect compared to wild-type cells, and cells were harvested at this point (**Table S5**). Cells were pelleted and resuspended in ddH_2_O at 300 OD_600_ units/mL. Low molecular weight RNA was isolated using the yeast hot-phenol method. Pelleted cells (300 OD_600_) were resuspended in 4mL RNA extraction buffer (0.1M NaOAc pH 5.2, 20mM EDTA pH 8.0, 1% SDS). Phenol (saturated with 100mM TrisCl pH 7.5) was added at a 1:1 ratio with resuspended cells and vortexed every 2.5 minutes for 30 seconds for a total of 20 minutes with incubation at 55°C in between vortexing. Cell debris was pelleted (5000 rpm for 6.5 minutes at 4°C) and the aqueous layer was transferred to a new tube and an equal volume of PCA was added, shaken to mix and centrifuged (5000xg, 20 minutes, 4°C). The PCA-extracted aqueous layer was transferred to a new tube and this step was repeated, followed by ethanol precipitation to pellet the total RNA. Pellets were resuspended in 900μL 1X TE pH 8.0 and 100μL 3M NaOAc pH 5.2 for a second ethanol precipitation. The final purified RNA pellets were resuspended in 500 μL ddH_2_O, and the total RNA was quantified by Nanodrop and stored at -80°C.

### Northern blotting

Purified RNA (2-10 μg) was resolved by electrophoresis on a 10% polyacrylamide, 8M urea gel after addition of 10X northern dye (50% glycerol, 0.3% bromophenol blue, 0.3% ethidium bromide). RNA was transferred to a Hybond N+ membrane (Amersham) using voltage transfer. RNA was crosslinked to the membrane using the Optimal Crosslink setting on a SpectroLinker XL-1000 UV crosslinker (Spectronics corporation). Membranes were visualized using either 5’- end radiolabeled or 5’-end biotin probes (**Table S2**).

For northerns with radiolabeled probes, membranes were pre-hybridized for three hours while rotating at 50°C in 25 mL pre-warmed pre-hybridization buffer containing 5x SSC (3 M NaCl and 0.3 M NaCitrate), 50% formamide, 5x Denhardt’s solution, 1% SDS, and 100 μg/mL salmon sperm DNA (denatured at 95°C for 5 minutes and chilled on ice before addition). Denhardt’s solution contained 2% w/v BSA fraction V, 2% w/v Ficoll 400, and 2% w/v polyvinylpyrrolidone. Pre-hybridization buffer was removed and replaced with pre-warmed hybridization buffer (5x SSC buffer, 50% formamide, 5x Denhardt’s solution, 1% SDS, 5% dextran sulfate) including ∼10 pmol of 5’-end labeled probe for incubation while rotating overnight at 50°C. Oligos were 5’-end radiolabeled at a final concentration of 4 μM with T4 PNK (NEB) and γ-^32^P-ATP after incubation for one hour at 37°C followed by enzyme inactivation at 72°C for 10 minutes. A BioGel P6 column (BioRad) removed excess labeled ATP. Each membrane was washed four times for 15 minutes each with low stringency wash buffer (2x SSC buffer and 0.05% SDS) at room temperature. Membranes were exposed for 2 hours and imaged on a Typhoon imager (Cytiva). To reprobe the same membrane for a different RNA species, the membrane was stripped with pre-boiled stripping buffer (1% SDS in ddH2O) at 85°C for 15 minutes while rotating, followed by storage in 1x TBE.

For northerns performed with 5’-biotin probes (protocol courtesy of KM McKenney, RP Connacher and AC Goldstroham, personal communication), membranes were washed in 2x SSC buffer for 15 minutes while rocking, and then pre-hybridized while rotating in pre-warmed Ultrahyb Ultrasensitive hybridization buffer (Thermo Scientific) at 42°C for 1 hour. 5 nM of the 5’-biotin DNA oligo was added to the pre-warmed Ultrahyb buffer and incubated while rotating overnight at 42°C. The membrane was washed two times with low stringency wash buffer (2x SSC, 0.1% SDS) and two times with high stringency wash buffer (0.1x SSC, 0.1% SDS) while rotating at 42°C for 15 minutes each. Biotin detection was performed using the Chemiluminescent Nucleic Acid Detection Module (Thermo Scientific 89880) and imaged with the Omega Lum G (GelCompany) with 10 second to 4 minute exposure, depending on the intensity of the bound probe. Reprobring was performed after stripping using the same conditions described above, although the pre-hybridization step could be omitted for re-probing with biotin labeled oligonucleotides. For both radiolabeled and biotin probes, hybridization buffer was reused up to 4 times, with addition of new biotinylated oligonucleotides for each northern experiment.

The intensity of hybridized RNA on each membrane was quantified using ImageQuant (Cytiva Life Sciences). Mature and precursor tRNA levels were normalized to 5S levels on each membrane (tRNA/5S). One strain on each membrane was then set to a value of 1 (as indicated in Figure legends) and the normalized intensity observed for all other strains normalized to the intensity of the same RNA in the standard strain, as indicated in each figure legend. Two-sample assuming equal variance t-tests were performed on triplicate measurements for each sample in order to determine statistical significance, as indicated on **Figure 2 and Figure 5**, and in **Table S1**.

## ACKNOWLEDGEMENTS

We would like to thank Eric Phizicky for providing yeast strains and tRNA overexpression plasmids used in this work and for valuable conversations during manuscript preparation. We also thank Anita Hopper and Regina Nostromo for yeast strains and for advice on genetic protocols and northern analysis. We thank Abi Hubacher for performing preliminary experiments. We thank Dmitri and Elena Kudryashov for assistance with chemiluminescent imaging experiments. This research was supported by NIH R01GM130135 (to J.E.J), and the OSU Center for RNA Biology Graduate Fellowship (to I.E.B).

## REFERENCES

Alexandrov A, Chernyakov I, Gu W, Hiley SL, Hughes TR, Grayhack EJ, Phizicky EM. 2006. Rapid tRNA decay can result from lack of nonessential modifications. Mol Cell 21: 87–96.

Bash-Imam Z, Therizols G, Vincent A, Laforets F, Polay Espinoza M, Pion N, Macari F, Pannequin J, David A, Saurin JC et al. 2017. Translational reprogramming of colorectal cancer cells induced by 5-fluorouracil through a miRNA-dependent mechanism. Oncotarget 8: 46219–46233.

Borchardt EK, Martinez NM, Gilbert WV. 2020. Regulation and Function of RNA Pseudouridylation in Human Cells. Annu Rev Genet 54: 309–336.

Callahan KP, Butler JS. 2010. TRAMP complex enhances RNA degradation by the nuclear exosome component Rrp6. J Biol Chem 285: 3540–3547.

Chan PP, Lowe TM. 2009. GtRNAdb: a database of transfer RNA genes detected in genomic sequence. Nucleic Acids Res 37: D93–97.

Chang JH, Jiao X, Chiba K, Oh C, Martin CE, Kiledjian M, Tong L. 2012. Dxo1 is a new type of eukaryotic enzyme with both decapping and 5’-3’ exoribonuclease activity. Nat Struct Mol Biol 19: 1011–1017.

Chatterjee K, Marshall WA, Hopper AK. 2022. Three tRNA nuclear exporters in S. cerevisiae: parallel pathways, preferences, and precision. Nucleic Acids Res 50: 10140–10152.

Chernyakov I, Whipple JM, Kotelawala L, Grayhack EJ, Phizicky EM. 2008. Degradation of several hypomodified mature tRNA species in Saccharomyces cerevisiae is mediated by Met22 and the 5’-3’ exonucleases Rat1 and Xrn1. Genes Dev 22: 1369–1380.

Cosentino C, Toivonen S, Diaz Villamil E, Atta M, Ravanat JL, Demine S, Schiavo AA, Pachera N, Deglasse JP, Jonas JC et al. 2018. Pancreatic beta-cell tRNA hypomethylation and fragmentation link TRMT10A deficiency with diabetes. Nucleic Acids Res 46: 10302–10318.

Davis DR. 1995. Stabilization of RNA stacking by pseudouridine. Nucleic Acids Res 23: 5020–5026.

Davis DR, Poulter CD. 1991. 1H-15N NMR studies of Escherichia coli tRNA(Phe) from hisT mutants: a structural role for pseudouridine. Biochemistry 30: 4223–4231.

Dewe JM, Whipple JM, Chernyakov I, Jaramillo LN, Phizicky EM. 2012. The yeast rapid tRNA decay pathway competes with elongation factor 1A for substrate tRNAs and acts on tRNAs lacking one or more of several modifications. RNA 18: 1886–1896.

Dichtl B, Stevens A, Tollervey D. 1997. Lithium toxicity in yeast is due to the inhibition of RNA processing enzymes. EMBO Journal 16: 7184–7195.

Ge J, Karijolich J, Zhai Y, Zheng J, Yu YT. 2017. 5-Fluorouracil Treatment Alters the Efficiency of Translational Recoding. Genes 8.

Giaever G, Chu AM, Ni L, Connelly C, Riles L, Veronneau S, Dow S, Lucau-Danila A, Anderson K, Andre B et al. 2002. Functional profiling of the Saccharomyces cerevisiae genome. Nature 418: 387–391.

Gillis D, Krishnamohan A, Yaacov B, Shaag A, Jackman JE, Elpeleg O. 2014. TRMT10A dysfunction is associated with abnormalities in glucose homeostasis, short stature and microcephaly. Journal of medical genetics 51: 581–586.

Gustavsson M, Ronne H. 2008. Evidence that tRNA modifying enzymes are important in vivo targets for 5-fluorouracil in yeast. RNA 14: 666–674.

Han L, Kon Y, Phizicky EM. 2015. Functional importance of Psi38 and Psi39 in distinct tRNAs, amplified for tRNAGln(UUG) by unexpected temperature sensitivity of the s2U modification in yeast. RNA 21: 188–201.

Holzmann J, Frank P, Loffler E, Bennett KL, Gerner C, Rossmanith W. 2008. RNase P without RNA: identification and functional reconstitution of the human mitochondrial tRNA processing enzyme. Cell 135: 462–474.

Hopper AK, Huang HY. 2015. Quality Control Pathways for Nucleus-Encoded Eukaryotic tRNA Biosynthesis and Subcellular Trafficking. Mol Cell Biol 35: 2052–2058.

Hoskins J, Butler JS. 2008. RNA-based 5-fluorouracil toxicity requires the pseudouridylation activity of Cbf5p. Genetics 179: 323–330.

Howell NW, Jackman JE. 2019. Impact of Chemical Modification on tRNA Function. eLS.

Howell NW, Jora M, Jepson BF, Limbach PA, Jackman JE. 2019. Distinct substrate specificities of the human tRNA methyltransferases TRMT10A and TRMT10B. RNA 25: 1366–1376.

Igoillo-Esteve M, Genin A, Lambert N, Desir J, Pirson I, Abdulkarim B, Simonis N, Drielsma A, Marselli L, Marchetti P et al. 2013. tRNA methyltransferase homolog gene TRMT10A mutation in young onset diabetes and primary microcephaly in humans. PLoS genetics 9: e1003888.

Jackman JE, Alfonzo JD. 2013. Transfer RNA modifications: nature’s combinatorial chemistry playground. Wiley interdisciplinary reviews RNA 4: 35–48.

Jackman JE, Montange RK, Malik HS, Phizicky EM. 2003. Identification of the yeast gene encoding the tRNA m1G methyltransferase responsible for modification at position 9. RNA 9: 574–585.

Juhling F, Morl M, Hartmann RK, Sprinzl M, Stadler PF, Putz J. 2009. tRNAdb 2009: compilation of tRNA sequences and tRNA genes. Nucleic Acids Res 37: D159–162.

Kadaba S, Krueger A, Trice T, Krecic AM, Hinnebusch AG, Anderson J. 2004. Nuclear surveillance and degradation of hypomodified initiator tRNAMet in S. cerevisiae. Genes Dev 18: 1227–1240.

Kempenaers M, Roovers M, Oudjama Y, Tkaczuk KL, Bujnicki JM, Droogmans L. 2010. New archaeal methyltransferases forming 1-methyladenosine or 1-methyladenosine and 1-methylguanosine at position 9 of tRNA. Nucleic Acids Res 38: 6533–6543.

Kotelawala L, Grayhack EJ, Phizicky EM. 2008. Identification of yeast tRNA Um(44) 2’-O-methyltransferase (Trm44) and demonstration of a Trm44 role in sustaining levels of specific tRNA(Ser) species. RNA 14: 158–169.

Kramer EB, Hopper AK. 2013. Retrograde transfer RNA nuclear import provides a new level of tRNA quality control in Saccharomyces cerevisiae. Proc Natl Acad Sci U S A 110: 21042–21047.

Kufel J, Tollervey D. 2003. 3’-processing of yeast tRNA(Trp) precedes 5’-processing. Rna 9: 202–208.

Lin H, Zhou X, Chen X, Huang K, Wu W, Fu J, Li Y, Polychronakos C, Dong GP. 2020. tRNA methyltransferase 10 homologue A (TRMT10A) mutation in a Chinese patient with diabetes, insulin resistance, intellectual deficiency and microcephaly. BMJ Open Diabetes Res Care 8.

Machnicka MA, Milanowska K, Osman Oglou O, Purta E, Kurkowska M, Olchowik A, Januszewski W, Kalinowski S, Dunin-Horkawicz S, Rother KM et al. 2013. MODOMICS: a database of RNA modification pathways--2013 update. Nucleic Acids Res 41: D262–267.

Murguia JR, Belles JM, Serrano R. 1996. The yeast HAL2 nucleotidase is an in vivo target of salt toxicity. J Biol Chem 271: 29029–29033.

Narayanan M, Ramsey K, Grebe T, Schrauwen I, Szelinger S, Huentelman M, Craig D, Narayanan V, Group CRR. 2015. Case Report: Compound heterozygous nonsense mutations in TRMT10A are associated with microcephaly, delayed development, and periventricular white matter hyperintensities. F1000Res 4: 912.

O’Connor JP, Peebles CL. 1991. In vivo pre-tRNA processing in Saccharomyces cerevisiae. Molecular & Cellular Biology 11: 425–439.

Payea MJ, Guy MP, Phizicky EM. 2015. Methodology for the High-Throughput Identification and Characterization of tRNA Variants That Are Substrates for a tRNA Decay Pathway. Methods Enzymol 560: 1–17.

Payea MJ, Hauke AC, De Zoysa T, Phizicky EM. 2020. Mutations in the anticodon stem of tRNA cause accumulation and Met22-dependent decay of pre-tRNA in yeast. RNA 26: 29–43.

Phizicky EM, Alfonzo JD. 2010. Do all modifications benefit all tRNAs? FEBS Lett 584: 265–271.

Phizicky EM, Hopper AK. 2023. The life and times of a tRNA. RNA 29: 898–957.

Siklar Z, Kontbay T, Colclough K, Patel KA, Berberoglu M. 2021. Expanding the Phenotype of TRMT10A Mutations: Case Report and a Review of the Existing Cases. J Clin Res Pediatr Endocrinol.

Stern E, Vivante A, Barel O, Levy-Shraga Y. 2022. TRMT10A Mutation in a Child with Diabetes, Short Stature, Microcephaly and Hypoplastic Kidneys. J Clin Res Pediatr Endocrinol 14: 227–232.

Strassler SE, Bowles IE, Dey D, Jackman JE, Conn GL. 2022. Tied up in knots: Untangling substrate recognition by the SPOUT methyltransferases. J Biol Chem 298: 102393.

Suzuki T. 2021. The expanding world of tRNA modifications and their disease relevance. Nat Rev Mol Cell Biol 22: 375–392.

Swinehart WE, Henderson JC, Jackman JE. 2013. Unexpected expansion of tRNA substrate recognition by the yeast m1G9 methyltransferase Trm10. RNA.

Tasak M, Phizicky EM. 2022. Initiator tRNA lacking 1-methyladenosine is targeted by the rapid tRNA decay pathway in evolutionarily distant yeast species. PLoS genetics 18: e1010215.

Torres AG, Batlle E, Ribas de Pouplana L. 2014. Role of tRNA modifications in human diseases. Trends in molecular medicine 20: 306–314.

Vilardo E, Amman F, Toth U, Kotter A, Helm M, Rossmanith W. 2020. Functional characterization of the human tRNA methyltransferases TRMT10A and TRMT10B. Nucleic Acids Res 48: 6157–6169.

Vilardo E, Nachbagauer C, Buzet A, Taschner A, Holzmann J, Rossmanith W. 2012. A subcomplex of human mitochondrial RNase P is a bifunctional methyltransferase--extensive moonlighting in mitochondrial tRNA biogenesis. Nucleic Acids Res 40: 11583–11593.

Whipple JM, Lane EA, Chernyakov I, D’Silva S, Phizicky EM. 2011. The yeast rapid tRNA decay pathway primarily monitors the structural integrity of the acceptor and T-stems of mature tRNA. Genes Dev 25: 1173–1184.

Yew TW, McCreight L, Colclough K, Ellard S, Pearson ER. 2016. tRNA methyltransferase homologue gene TRMT10A mutation in young adult-onset diabetes with intellectual disability, microcephaly and epilepsy. Diabetic medicine : a journal of the British Diabetic Association 33: e21–25.

Yun JS, Yoon JH, Choi YJ, Son YJ, Kim S, Tong L, Chang JH. 2018. Molecular mechanism for the inhibition of DXO by adenosine 3’,5’-bisphosphate. Biochemical and biophysical research communications 504: 89–95.

Zung A, Kori M, Burundukov E, Ben-Yosef T, Tatoor Y, Granot E. 2015. Homozygous deletion of TRMT10A as part of a contiguous gene deletion in a syndrome of failure to thrive, delayed puberty, intellectual disability and diabetes mellitus. Am J Med Genet A 167A: 3167–3173.

